# A New approximate matching compression algorithm for DNA sequences

**DOI:** 10.1101/853358

**Authors:** J.M. Lázaro-Guevara, K.M. Garrido

## Abstract

1.

Undeveloped countries like Guatemala, where access to high-speed internet connections is limited, downloading and sharing Biological information of thousands of Mega Bits is a huge problem for the beginning and development of Bioinformatics. Based on that information is an urgent necessity to find a better way to share this biological data. There is when the compression algorithms become relevant. With all this information in mind, born the idea of creating a new algorithm using redundancy and approximate selection.

**Methods:** Using the probability given by the transition matrix of the three-word tuple and relative frequencies. Calculating the relative and total frequencies given by the permutation formula (nr) and compressing 6 bits of information into 1 implementing the ASCII table code (0…255 characters, 28), using clusters of 102 DNA bases compacted into 17 string sequences. For decompressing, the inverse process must be done, except that the triplets must be selected randomly (or use a matrix dictionary, 4102).

**Conclusion:** The compression algorithm has a better compression ratio than LZW and Huffman’s algorithm. However, the time needed for decompressing makes this algorithm incompatible for massive data. The functionality as MD5sum need more research but is a promising helpful tool for DNA checking.

## 2. Background

Undeveloped countries like Guatemala, where the access to high speed internet connections is limited downloading and sharing Biological information of thousands of Mega Bits is a huge problem for the beginning and development of Bioinformatics.

Based on that information is an urgent necessity to find a better way to share this biological data. There are two different ways to solve this problem, the first one improving the velocity at which the data is transferred (improving the internet connection), however this is not affordable in a country like Guatemala, the second way is to decrease the amount of data transferred.[1][2]

To accomplish this goal on reducing the data, the most common form is compressing the information, and for compressing the information there are several ways to do it.

But this compression forms depends on the kind of information or data to be compressed, the most reliable methods like Huffman Algorithm, cannot be used in the DNA compression because this methods based on relative frequencies can involve some data lost (~3%)[1]. A second way is using a more reliable method like **LZW (Lempel-Ziv-Welch)** where there is no lost on data but need the use of license to generate the compressed code based on patents, and involves the use of a matrix dictionary for each code compressed[2][3].

With all this information in mind, born the idea of creating a new algorithm based on redundancy and approximate probability given by the transition matrix of the three word tuple and relative frequencies [3][4](figure 1).

**FIGURE 1.**
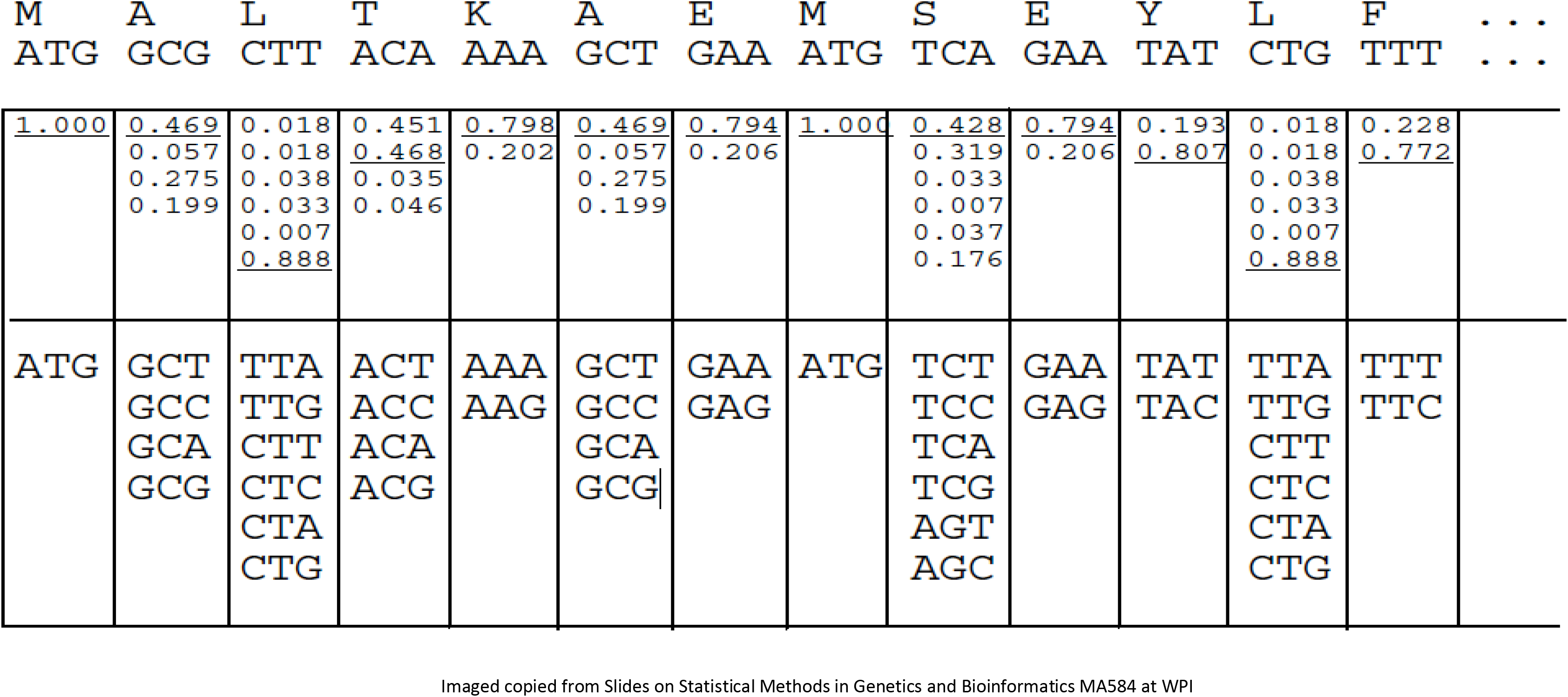

## 3. Methods

### 3.1 Compressing Method

First Step: Get a DNA sequence, and split it in subsequence of 102 bases

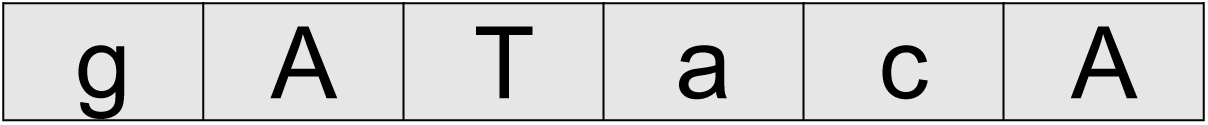

Second Step: transform the bases to a numeric correspondent (a=1, c=2, g=3, t=4).

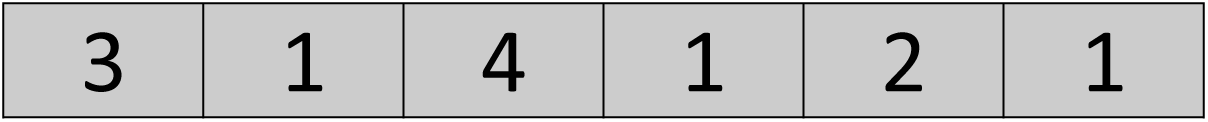

Third Step: It is necessary to divide in 102 bases because is needed a number than can be divided by 6 and 3, and also superior a 100 to improve the compression ratio, based and that 1 character will represent a 6 DNA bases.

a. 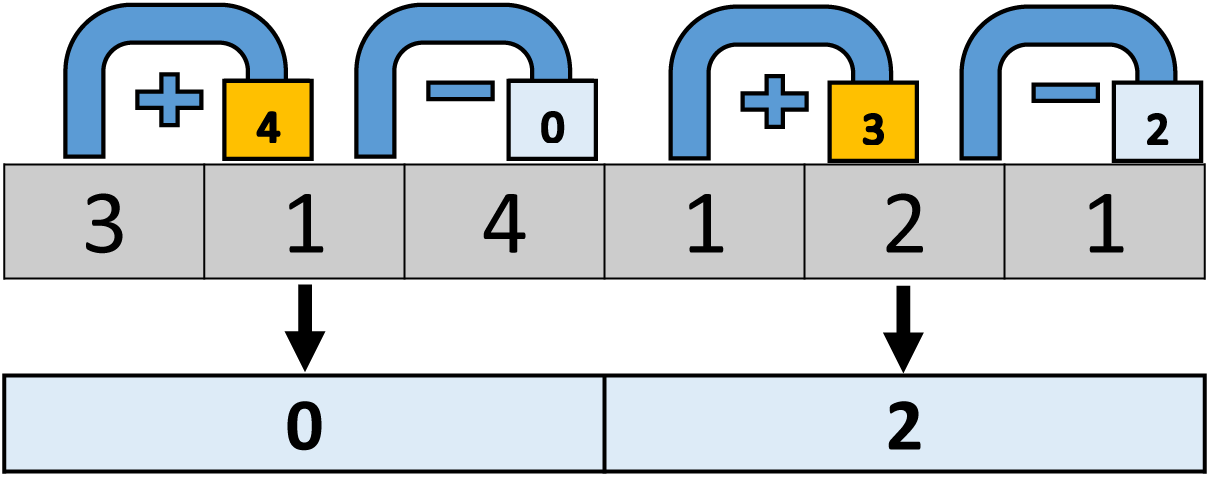
b. Using permutation **n × n × …(r times) = n^r^,** r times, we have 4 possibilities (a,c,g,t), and selecting 3 each time, based on the process of adding the first one and subtracting to that sum the third value (eg. (3+1)−4=0), we only have 7 possible values that will be from **−2 to 7**. This model of adding and subtracting was used to obtain an array of 10 elements in a normal distribution (1/64,3/64,6/64,10/64,12/64) (table 1).

**TABLE I.**
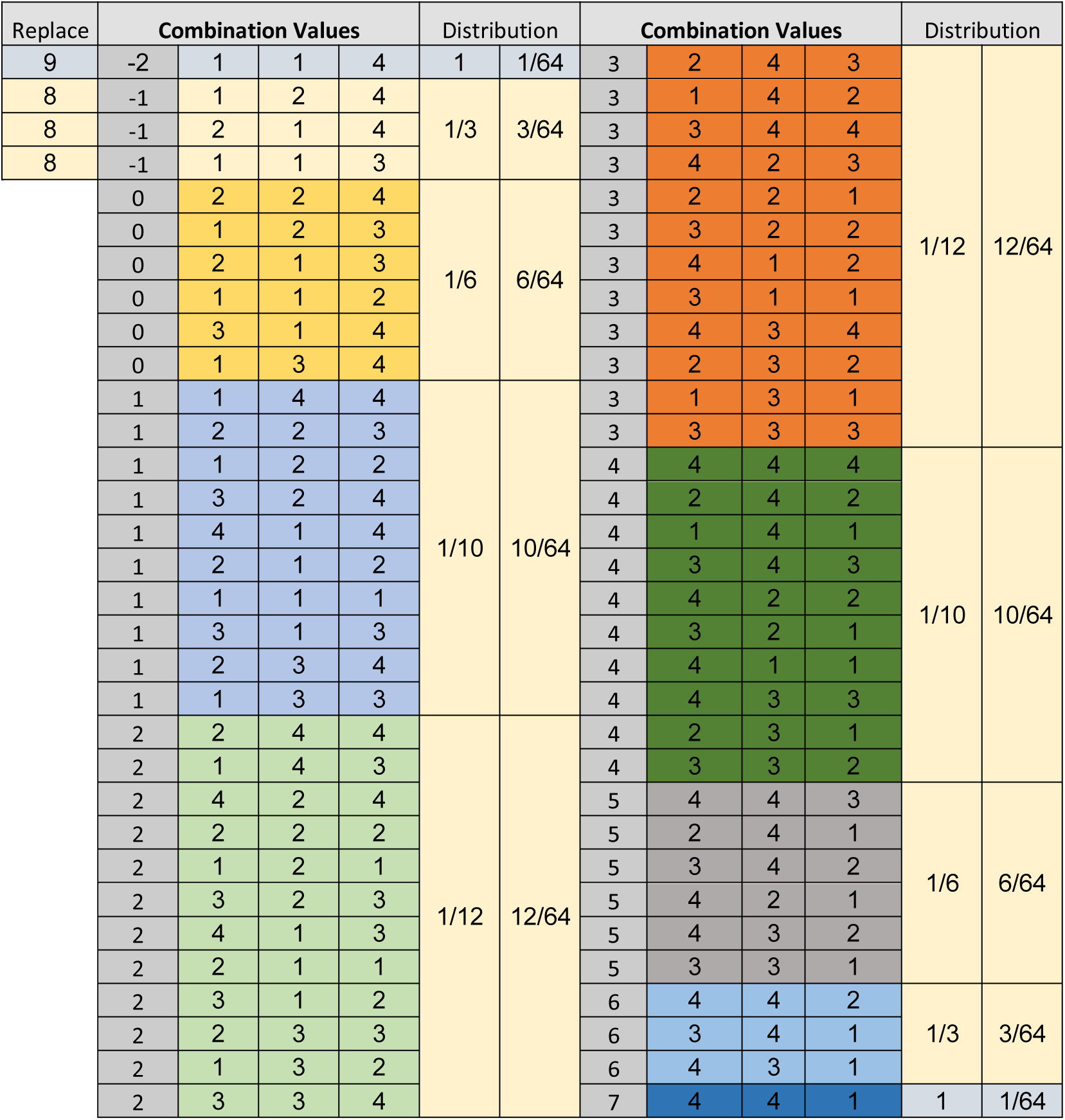
64 permutations in Triplets for DNA encoding and d ecoding (Numerical representation of DNA Triplets)
c. It is necessary to group 64 possible combination of values in only 10 combinations to use the ASCII mode (range from 0 to 255), in table I it is possible to observe that exist two negative values (−2,−1), and it is necessary to replace them in the second value by the (8 = −1, and 9 = −2), to use the concatenation of two characters as one number.
d. After obtaining 2 characters and concatenating it, it is necessary to add 63 to this value, to avoid characters used by the core system. The range of values after concatenation will go from −29 to 79. So after adding 63 to this values the range in ASCII will be from 34 to 142. Table II.

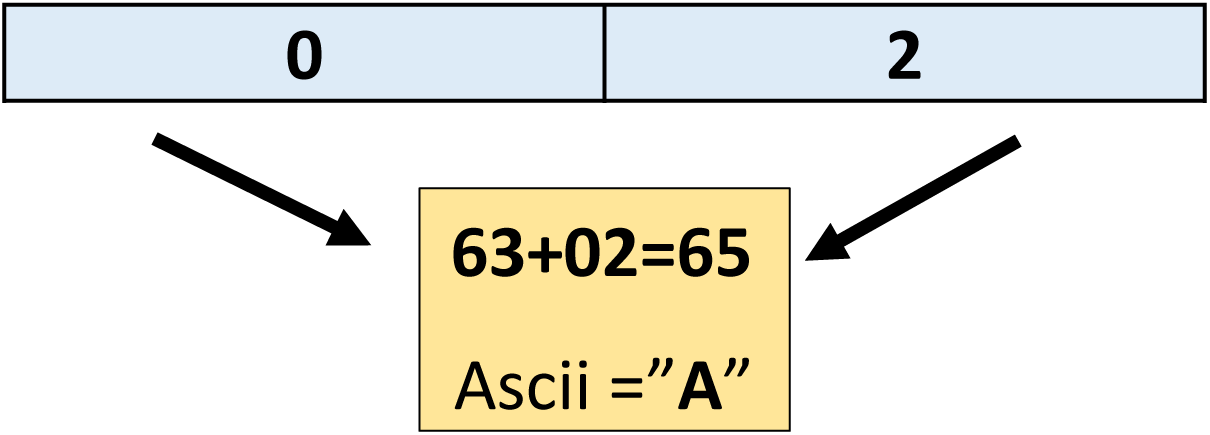

**TABLE II.**
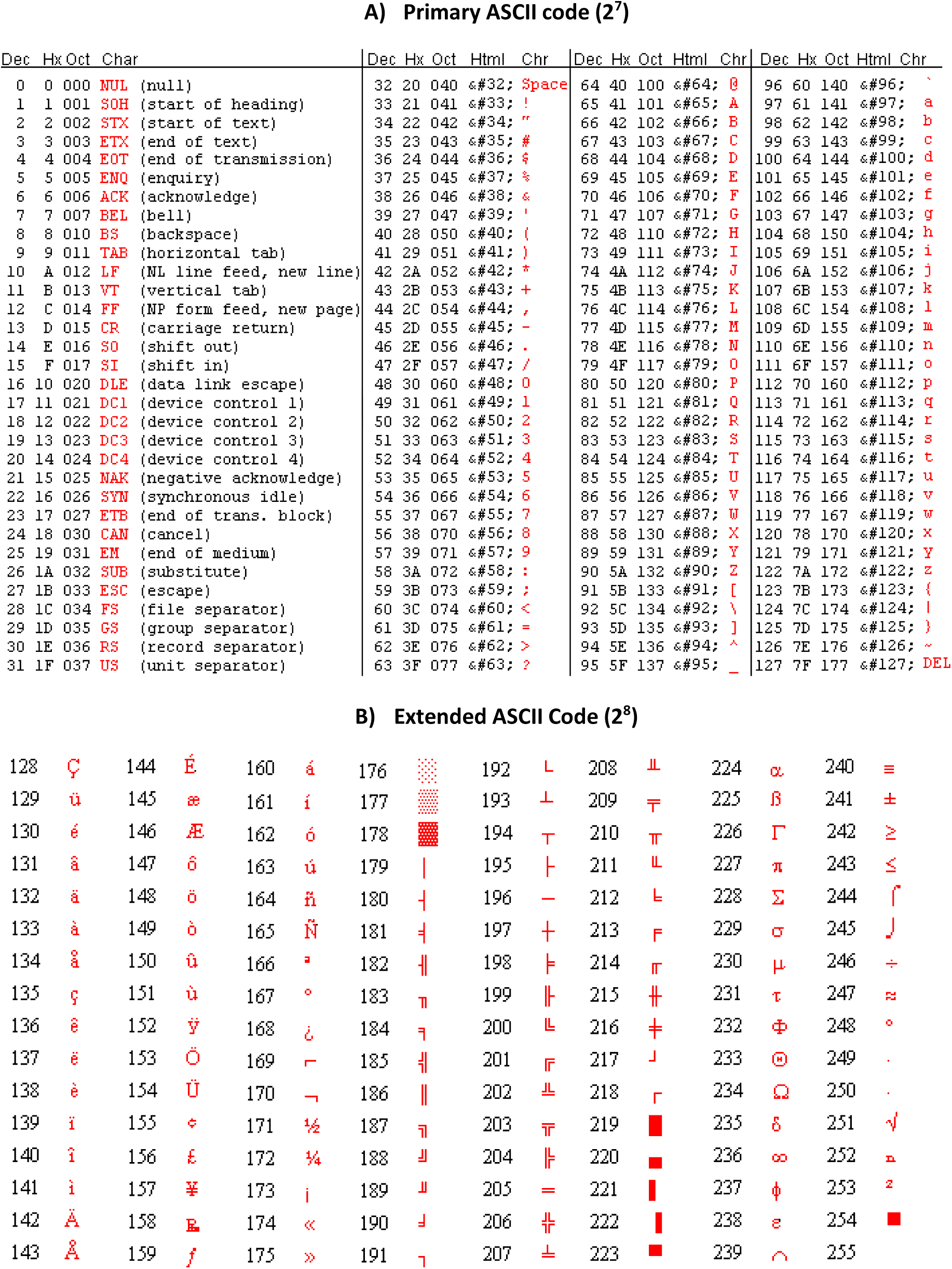
**ASCII SEQUENCE FOR CHAR AND HEXADECIMAL COMPRESION** Example 1: **DNA code: (102 characters)** a t a a c c a c a a a g a g c a g g t a a c g a g c c c g c a g c t t g t a a g c g c g a g c t a t t a a c a a t c g t t c c a c t t g g c a a c g c t a c c a g g a t g t t a a c a a a g t c c a t a g g **Compressed Version: (17 characters)** h[Tk’XihI*oi’?]N4
e. Checking numbers 1 and 2: This numbers allow to calculate the original 102 bases compacted into 17 characters from 34 numbers. Number 1 is obtained by adding and subtracting the product of the 34 calculated number. (eg. (3+1−4)−(1+2−1)…. +….−). In case of example 1 is **11.** Number 2 is obtained by a similar process using multiplication and subtracting process calculated number of the 34 by the formula (2*(1st*3rd))−(3*2nd) (eg. (2(3*4)−(3*1))−(2(1*1)−(3*2)…. +….−). **In case of example 1 is −3.** Compacted code of example 1: **h[Tk’XihI*oi’?]N4!11!−3**
f. Line 1 header. This is the first line of the compacted code and contains the absolute frequency of each one of the 64 possible combinations for future checking in the decompressing process. **(eg. 62-95-94-59-06-34-59-46-27-62-66-60-59-96-43-63-36-16-62-56-43-64-26-44)**

### 3.2 Decompressing Method

a. Read Line 1, introduce the data into a linear vector with relative frequencies of each triplet for further comparison against the decompressed code and the original code, and improving the velocity of the change based on the change in the total amount of probability of each possible term.
b. Obtain the compacted code (e.g **h[Tk’XihI*oi’?]N4!11!-3**), split into three different terms using the symbol “**!”**, as separation indicator. **Term 1:** 17 character code. **Term 2:** number 1 and **Term 3:** number 2.
c. Using the reverse method of part d in compression method, convert the ascii code into a two digit value (eg. Asci = “A”, number from asci = 65), subtracts **63** to that number (eg. 65 - 63=02), using the modulus and the integer division separate into 2 numbers (02 = 0, 2). If number two is 8 or 9 change into the negative portion (eg. **− 29 = − 2, − 2**).
d. From the 17 character code, after changing in the 2 digits way, it will be converted into a 34 digits code.
e. Now is necessary to change 34 digits into 102 characters, but it is no possible to use the inverted process on step b on the codifying part because each number has may possibilities from 3 to 12 possible options, only numbers −2 and 7, has one direct transcription, and number −2 and 7, where selected based on the most common triplet of bases repeated in 1000 trials of codifying sequences.
f. Because there are 102 possible DNA bases in this code and there are 4 possible letter for sequence and the order it is important, it is necessary to calculate the permutation **4 P 102**, that it is equivalent to count 34 digits and 64 possible values obtained from the triplets, **64 P 34**. But this value is so huge (2.57 × 10E61), but it is limited as shown in TABLE III, by the number of probabilities from 3 to 12, in 8 possible numbers.

**TABLE III.** Sequence of DNA of Example 1, possible results on checking numbers 1 and 2 after permutations. For that reason is necessary the checking numbers 1 and 2, there is only one sequence of 34digits (17 characters) for each pair of checking numbers (look at probability table at the bottom of TABLE III, figure 1).

|  | 4 | 1 | 2 | -1 | 2 | 1 | 4 | 4 | 3 | 3 | 2 | 5 | 4 | 2 | 4 | 1 | 1 | 0 | -2 | 1 | 4 | -1 | 4 | 2 | 3 | 3 | 0 | 0 | 3 | 0 | 1 | 5 | -1 | 1 |
| --- | --- | --- | --- | --- | --- | --- | --- | --- | --- | --- | --- | --- | --- | --- | --- | --- | --- | --- | --- | --- | --- | --- | --- | --- | --- | --- | --- | --- | --- | --- | --- | --- | --- | --- |
| 1 | 444 | 144 | 244 | 124 | 244 | 144 | 444 | 444 | 243 | 243 | 244 | 443 | 444 | 244 | 444 | 144 | 144 | 224 | 114 | 144 | 444 | 124 | 444 | 244 | 243 | 243 | 224 | 224 | 243 | 224 | 144 | 443 | 124 | 144 |
| 2 | 242 | 223 | 143 | 214 | 143 | 223 | 242 | 242 | 142 | 142 | 143 | 241 | 242 | 143 | 242 | 223 | 223 | 123 |  | 223 | 242 | 214 | 242 | 143 | 142 | 142 | 123 | 123 | 142 | 123 | 223 | 241 | 214 | 223 |
| 3 | 141 | 122 | 424 | 113 | 424 | 122 | 141 | 141 | 344 | 344 | 424 | 342 | 141 | 424 | 141 | 122 | 122 | 213 |  | 122 | 141 | 113 | 141 | 424 | 344 | 344 | 213 | 213 | 344 | 213 | 122 | 342 | 113 | 122 |
| 4 | 343 | 324 | 222 |  | 222 | 324 | 343 | 343 | 423 | 423 | 222 | 421 | 343 | 222 | 343 | 324 | 324 | 112 |  | 324 | 343 |  | 343 | 222 | 423 | 423 | 112 | 112 | 423 | 112 | 324 | 421 |  | 324 |
| 5 | 422 | 414 | 121 |  | 121 | 414 | 422 | 422 | 221 | 221 | 121 | 432 | 422 | 121 | 422 | 414 | 414 | 314 |  | 414 | 422 |  | 422 | 121 | 221 | 221 | 314 | 314 | 221 | 314 | 414 | 432 |  | 414 |
| 6 | 321 | 212 | 323 |  | 323 | 212 | 321 | 321 | 322 | 322 | 323 | 331 | 321 | 323 | 321 | 212 | 212 | 134 |  | 212 | 321 |  | 321 | 323 | 322 | 322 | 134 | 134 | 322 | 134 | 212 | 331 |  | 212 |
| 7 | 411 | 111 | 413 |  | 413 | 111 | 411 | 411 | 412 | 412 | 413 |  | 411 | 413 | 411 | 111 | 111 |  |  | 111 | 411 |  | 411 | 413 | 412 | 412 |  |  | 412 |  | 111 |  |  | 111 |
| 8 | 433 | 313 | 211 |  | 211 | 313 | 433 | 433 | 311 | 311 | 211 |  | 433 | 211 | 433 | 313 | 313 |  |  | 313 | 433 |  | 433 | 211 | 311 | 311 |  |  | 311 |  | 313 |  |  | 313 |
| 9 | 231 | 234 | 312 |  | 312 | 234 | 231 | 231 | 434 | 434 | 312 |  | 231 | 312 | 231 | 234 | 234 |  |  | 234 | 231 |  | 231 | 312 | 434 | 434 |  |  | 434 |  | 234 |  |  | 234 |
| 10 | 332 | 133 | 233 |  | 233 | 133 | 332 | 332 | 232 | 232 | 233 |  | 332 | 233 | 332 | 133 | 133 |  |  | 133 | 332 |  | 332 | 233 | 232 | 232 |  |  | 232 |  | 133 |  |  | 133 |
| 11 |  |  | 132 |  | 132 |  |  |  | 131 | 131 | 132 |  |  | 132 |  |  |  |  |  |  |  |  |  | 132 | 131 | 131 |  |  | 131 |  |  |  |  |  |
| 12 |  |  | 334 |  | 334 |  |  |  | 333 | 333 | 334 |  |  | 334 |  |  |  |  |  |  |  |  |  | 334 | 333 | 333 |  |  | 333 |  |  |  |  |  |

### 3.3 Measuring Accuracy on decompression and data loss

For obtaining the original sequence of DNA it is necessary to obtain the permutations, and compare it to the checking numbers, the problem is that this take a lot of time so the best solution is to import a common Tables Comparing Dictionary, the disadvantage of this solution is the big size of the dictionary.

The measuring of accuracy is related to the number of cycles permuted, to obtain the original DNA sequence is necessary compare checking numbers 1 and 2, until obtaining the original sequence, more the numbers of permutation higher the probability on obtaining the 102 original DNA bases per permutation. (TABLE IV)

**TABLE IV.**
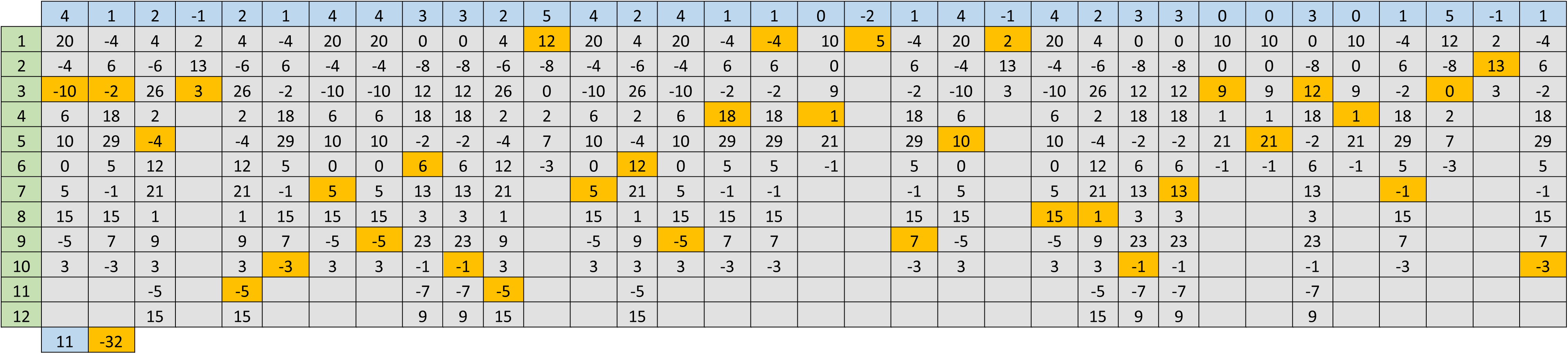
Sequence of DNA of Example 1, checking numbers 1 and 2.

The accuracy will be measured comparing simultaneously the sequences (Compressed = original), and measuring the differences or sections with discrepancy it is obtained the accuracy proportion (e.g)

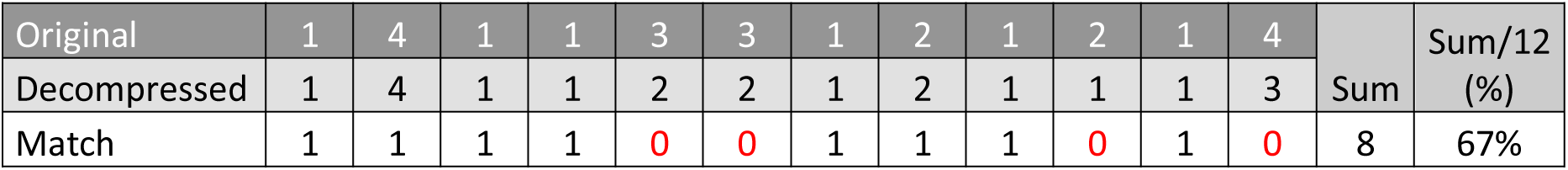

## 4. Results

After several trials, and running multiple simulations in R (Statistical Programming Language), on how this compression and decompression system works. (Figure 2, figure 3) It is obtained a compression ratio based on data weight in kilobytes (Kb), using a 10,000 bases (A, C, G, T) section of DNA extracted from NC_000913.1 (E.coli) with read. Genbank function.

**FIGURE 2.**
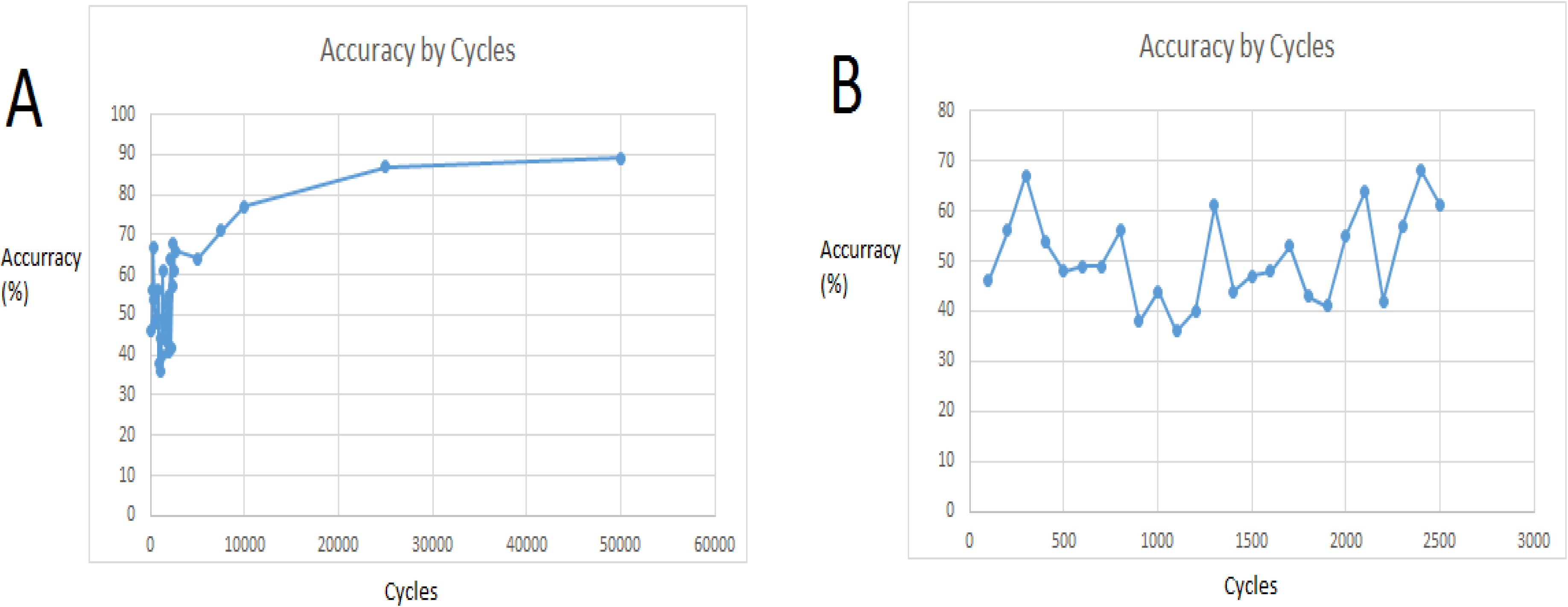

**FIGURE 3.**
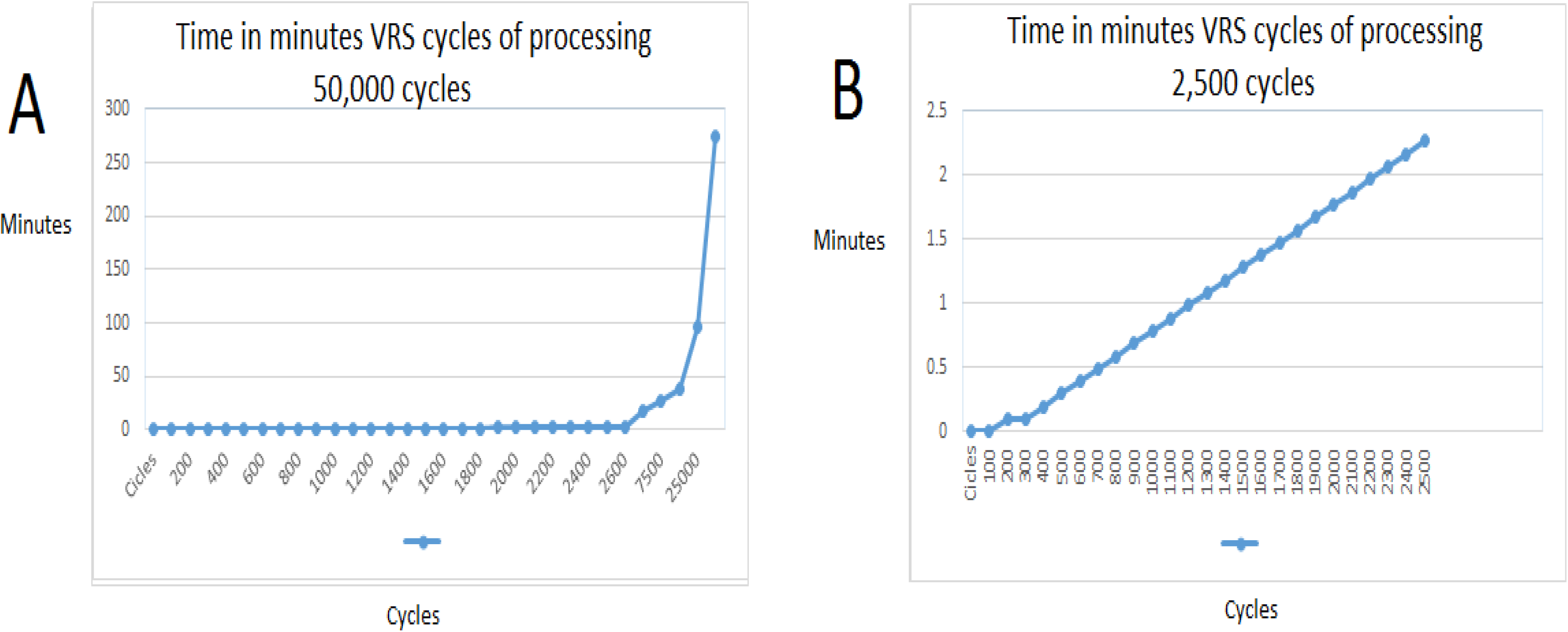

The data extracted and printed in a text file (.TXT), uses 20,100 bytes, the data compacted in the same archive system uses only 2,567.

The definition of the compression ratio is (|O|/2| I|), where |I| is number of bases in the input DNA sequence and |O| is the length (number of bits) of the output sequence, 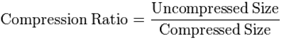, **the general compression ratio is 8:1** (7.84), the **inline code compression ratio is 5:1** (102 DNA bases are equivalents to 17 ASCII codes plus 2 checking numbers). The compression ratios, as well as the multiple permutation system, are presented in Figure 1.

The compressing and decompressing time is related to the number of cycles (~102 permutations), using a quad-core 2.3 Ghz processor as standard, it takes **38 milliseconds** to complete one cycle, but because it is using a iteration systems (for and while commands), taking on count the RAM and processor bus transfer speed is necessary to add 15% on delay each 1000 cycles.

The accuracy on decoding is related directly to the number of iterations and permutations (figure 2), whit only 100 to 2,500 accuracy varies from 50 to 70%, but this accuracy is counted only in a line of decoding (102 DNA bases), based on time of processing, after 7,500 cycles accuracy increase in exponential, in range from 80 to 91%, reaching the limit in cycles around 50,000, because Random Access Memory limiting, so it is unknown if the accuracy will reach 100% in the estimated 450,000 cycles (RAM memory insufficient for this calculation on normal PC 8 Gigabytes).

After running a sequence of 1020 DNA bases, using 5000 cycles, the analysis took 9.5 hours, and accuracy obtained was 63%. A combination dictionary for improving the speed and accuracy was created with 120,000 permutations and speed increase by 5%, for creating the complete dictionary with all possible permutations (2.57 × 10E61), certainly will improve accuracy and speed of decoding, but was not possible to create it with the R programming language, based on the RAM size.

## 5. Discussion

This algorithm have demonstrated that the general compression ratio obtained (8:1) is better that traditional compression codes (eg LZW, Huffmans, Biocompress, Gencompress, FASTA), but the time needed for decompression makes no worthy the usage of this compression method by using multiple permutations, it will improve using a dictionary of sequences preinstalled in the coding and decoding software.

It was not possible to probe the complete accuracy, assuring no data lose in the decompression system after 50,000 cycles accuracy only reach 86%, adding some modifications like improving the random sampling permutations it was able to obtain 91%. For that reason is necessary to do more changes on the code and try to reach 100% accuracy on selected sampling instead of random sampling on a better equipment with more processor speed and RAM.

Besides the time consumption for decompressing, this algorithm can also be used in the compression form as a checksum like the MD5sum system for DNA codes. MD5 algorithm is the mainstream for the cryptographic check and file checkup, the hash function when MD5 is applied returns a unique check number that allow knowing that this file is uncorrupted, this is an important problem in compressing method when data lose exist, and is necessary to be sure of that DNA is not corrupted.

So it is necessary to continue in researching for improving it.

## 6. Conclusion

- The compression algorithm has better compression ratio than LZW and huffmans algorithm.
- The time need for decompressing make not useful this algorithm for huge amount of data.
- The functionality as MD5sum need more research, but is a promising useful tool for DNA checking.

---

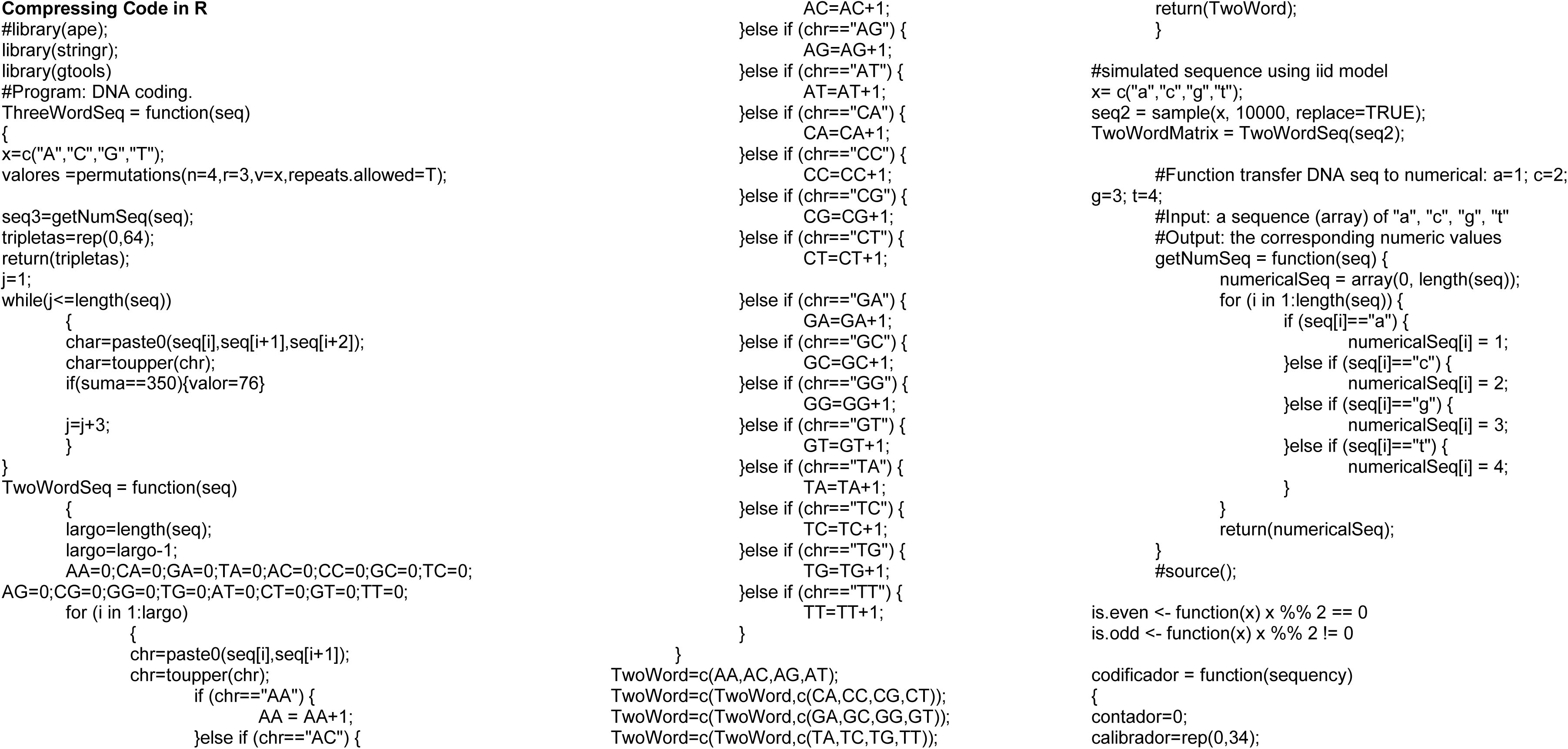

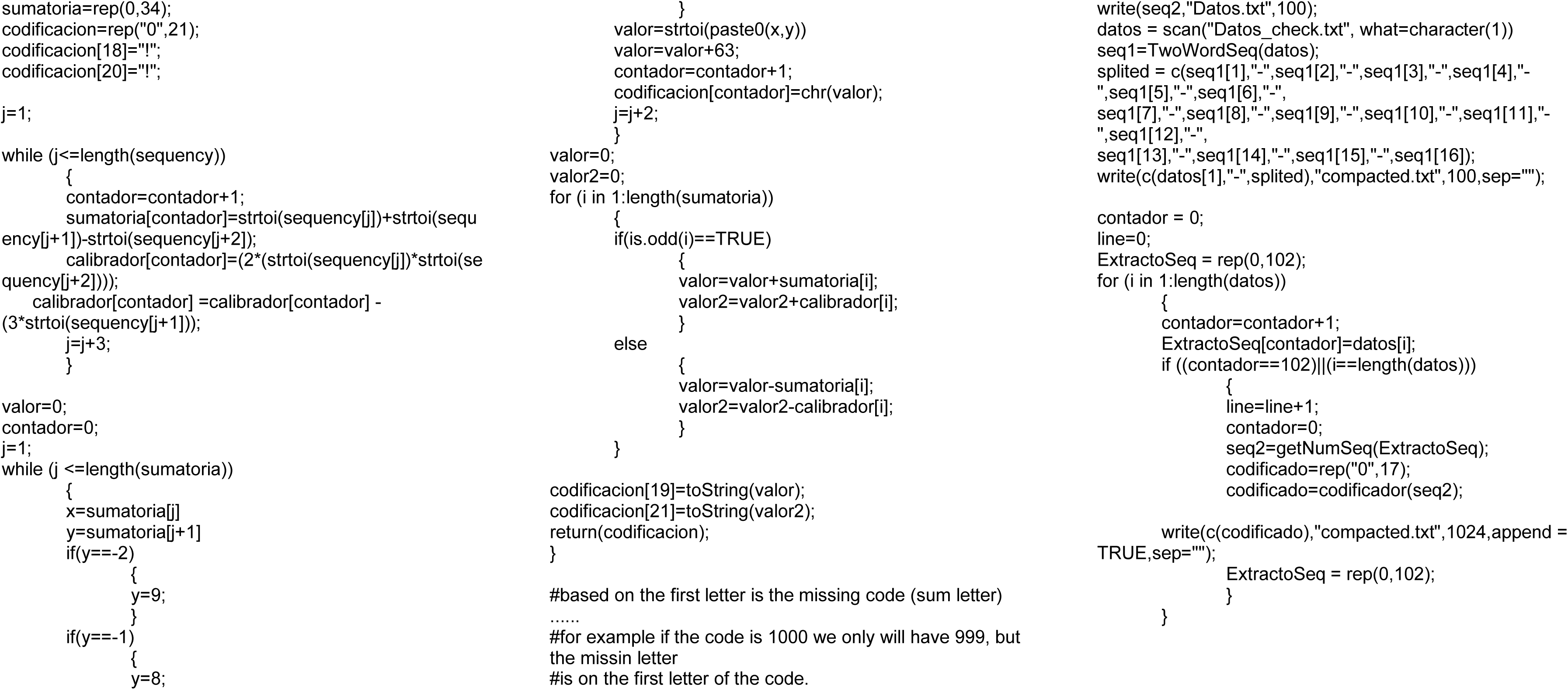

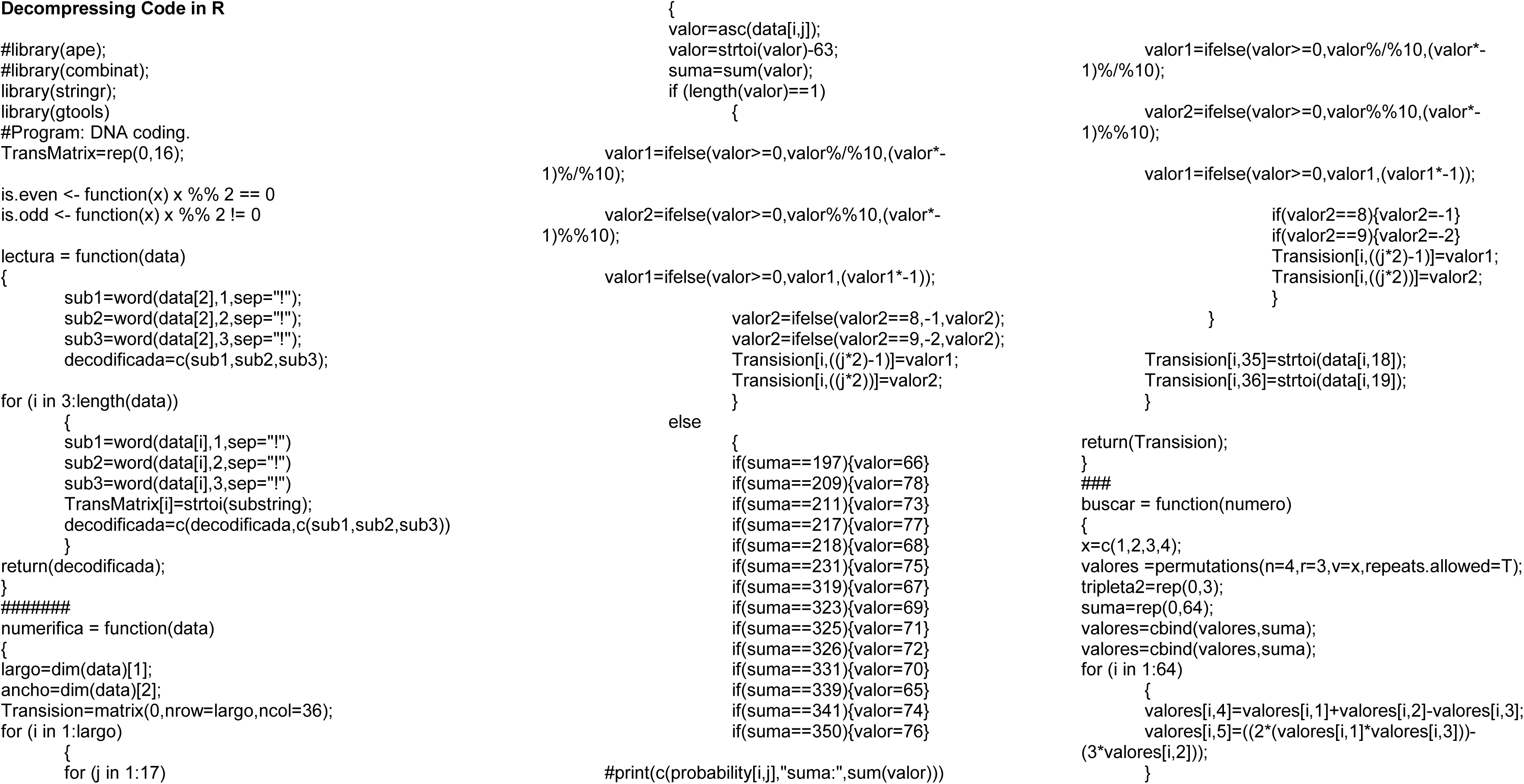

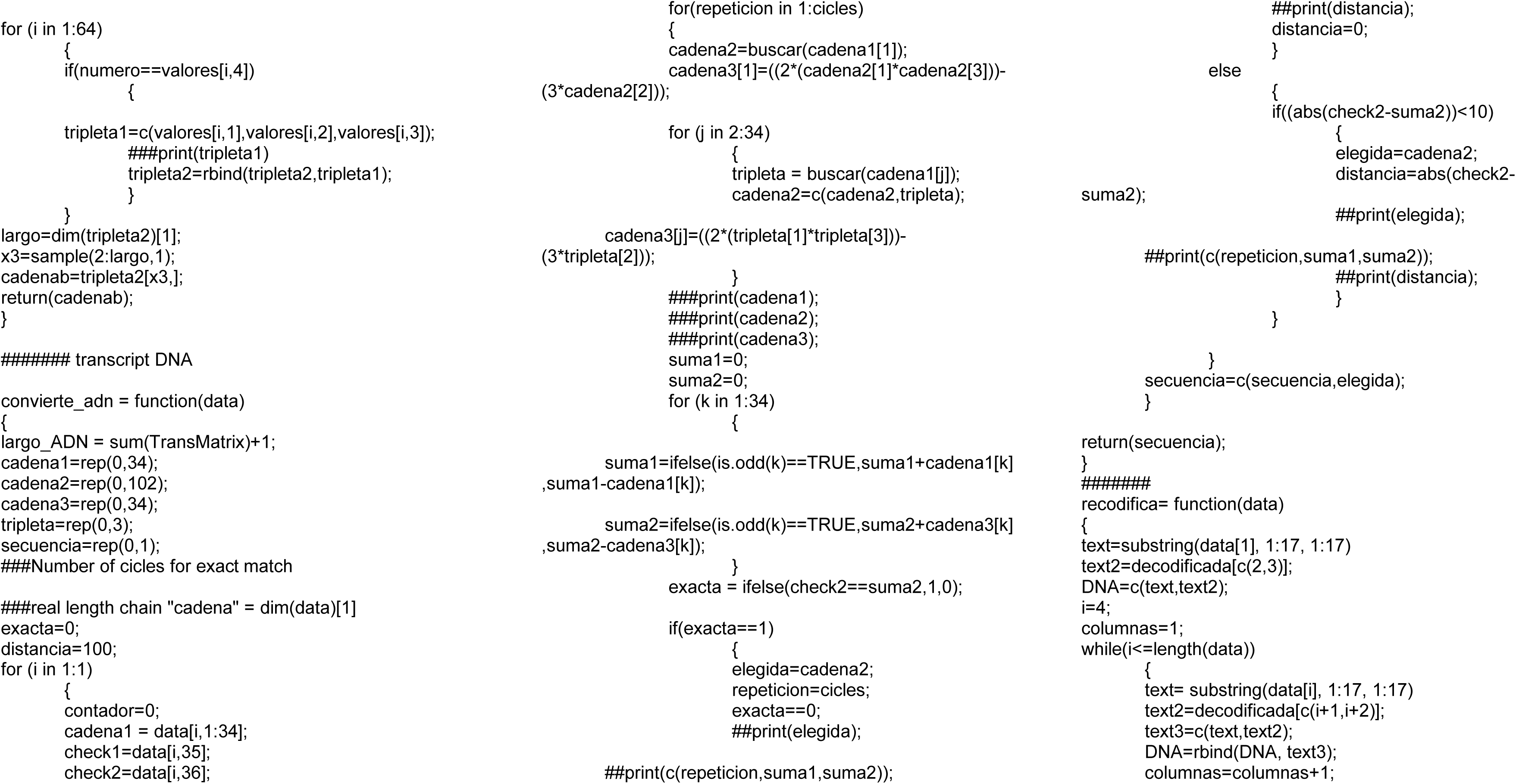

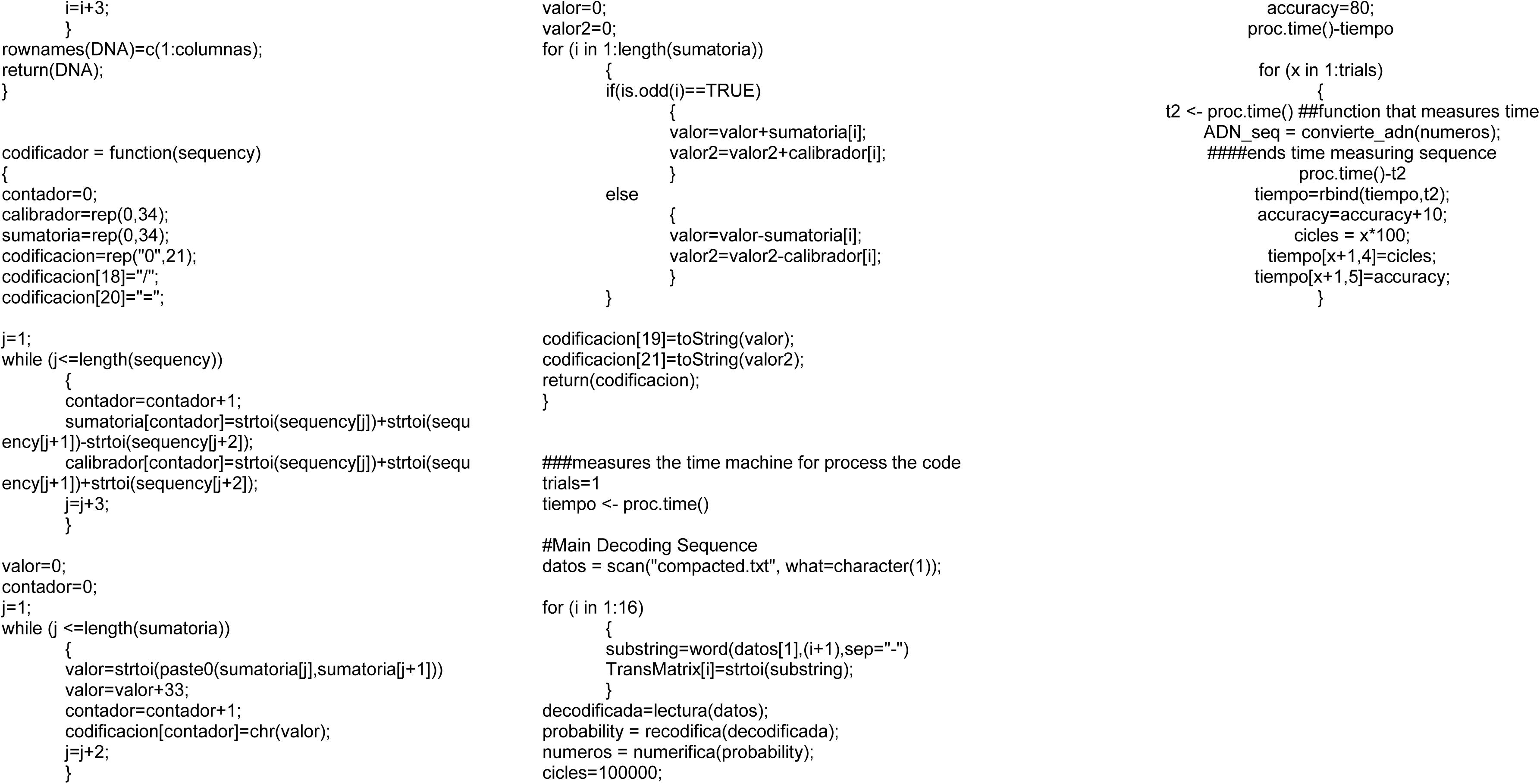

